# Microscale light management and inherent optical properties of intact corals studied with optical coherence tomography

**DOI:** 10.1101/376723

**Authors:** Daniel Wangpraseurt, Steven Jacques, Niclas Lyndby, Jacob Boiesen Holm, Christine Ferrier Pages, Michael Kühl

## Abstract

Coral reefs are highly productive photosynthetic systems and coral optics studies suggest that such high efficiency is due to optimised light scattering by coral tissue and skeleton. Here, we characterise the inherent optical properties, i.e., the scattering coefficient, ***μ**_s_*, and the anisotropy of scattering, *g*, of 8 intact coral species using optical coherence tomography (OCT). Specifically, we describe light scattering by coral skeletons, coenoarc tissues, polyp tentacles and areas covered by fluorescent pigments (FP). Our results reveal that light scattering between coral species ranges from ***μ**_s_* = 3 mm^−1^ (*Stylophora pistillata*) to *μs*= 25 mm^−1^ (*Echinopora lamelosa*). For *Platygyra pini, **μ**_s_* was 10-fold higher for tissue vs skeleton, while in other corals (e.g. *Hydnophora pilosa*) no difference was found between tissue and skeletal scattering. Tissue scattering was 3-fold enhanced in coenosarc tissues (***μ**_s_* = 24.6 mm^−1^) vs polyp tentacles (***μ**_s_* = 8.3 mm^−1^) in *Turbinaria reniformis*. FP scattering was almost isotropic when FP were organized in granule chromatophores (*g*=0.34) but was forward directed when FP were distributed diffusely in the tissue (*g*=0.96). Our study provides detailed measurements of coral scattering and establishes a rapid approach for characterising optical properties of photosynthetic soft tissues via OCT *in vivo*.

## Introduction

The form and function of an organism represents a design solution to the problems posed by a multitude of environmental parameters. The evolutionary design of terrestrial plants has been studied over decades, providing evidence on the prime role of irradiance exposure, hydration and mechanical stability in driving plant morphology on cellular to canopy scales [1, 2]. On the scale of a plant leaf, studies showed that epidermal cells act similar to a lens and can focus incident radiation [3]. Light focused by epidermal cells can then be channelled deep into the plant leaf via lossy scattering between air-filled vacuoles and the palisade layer (i.e. a porous matrix) [4]. Such light propagating mechanism is more pronounced in sun adapted leaves that are in need of effectively distributing excess irradiance [4] compared to shade adapted leaves which did not have such well-defined palisade mesophyll and instead developed a spongy mesophyll, which scatters less light [5]. It is also known that some succulents that are exposed to high light (e.g. *Dudleya brittonii*) can develop wax cuticles that effectively attenuate UV light and could thus act photoprotective [6]. Thus for terrestrial plants, the plant leaf is highly specialised in optimising irradiance exposure for chloroplast photosynthesis [3].

On tropical coral reefs, corals have evolved as highly efficient photosynthetic organisms [7]. Similar to plants, corals can be subject to variable irradiance regimes. Shallow water corals are exposed to excess irradiance, reaching about 2000 μmol photons m^−2^ s^−1^ during mid-day sun and low tide [8]. In contrast, deep water corals can live photosynthetically under mesophotic conditions leaving only about 3% of surface irradiance at a depth of over >100 m [9]. While it is well-known that the coral animal’s symbiotic microalgae (*Symbiodinium* sp.) adjusts pigment density (specifically chlorophyll *a*) to optimise light absorption for photosynthesis [10] there is now mounting evidence that the light scattering properties of coral skeletons and tissues also play a central role in light propagation and coral photosynthesis [11–16].

Studies with fiber-optic light microsensors showed that the coral tissue surface scalar irradiance (i.e. the total quantum flux in a point incident from all directions) can be about twice of the incident downwelling irradiance [11]. Similar to plant tissues, coral tissues are suggested to be strongly light scattering matrices [15]. Light scattering affects the photon pathlength and residence time in a given tissue layer and thus the chance for light absorption and subsequent O_2_ evolution or heat dissipation. Monte Carlo simulations (i.e. probabilistic light propagation models [17]) were developed to quantify the scattering probability of coral tissue and skeletons [16]. It was found that the scattering coefficient, ***μ**_s_* [mm^−1^], i.e., the ability to scatter photons over a certain distance, of the massive coral *Favites abdita*, was about 3 times higher in the tissue than in the underlying skeleton, indicating that light is effectively homogenized by the tissue and then propagated by the skeleton [16]. Other studies investigated light scattering from a museum collection of coral skeletons and showed that scattering is highly variable between coral skeletons [14, 18]. Differences in light scattering were related to coral bleaching susceptibility [14] and corals with low light scattering ability appeared more susceptible to coral bleaching [19]. Despite a few studies that have extracted the scattering coefficient of coral skeletons [14, 16, 19, 20] the current data on intact corals and coral tissues is very limited [16].

Characterisation of optical properties of soft tissues in their intact state is difficult given that many techniques rely on tissue preparation that can result in changes in tissue structure and hydration with the potential to alter tissue optical properties [21]. In the field of biomedical tissue optics, several approaches have been developed for extracting optical properties of intact tissues. A common technique is diffuse optical spectroscopy, where the lateral attenuation of diffuse reflectance is measured with fibers placed at several radial distances [21]. However, a key requirement for using diffusion theory is that the collected light is entirely diffuse, i.e., has lost directionality. Under the assumption of diffuse light, the diffuse scattering coefficient can be extracted as ***μ**_s_*’=***μ**_s_*(1-*g*), where *g* is the so-called anisotropy factor. The *g* value [dimensionless] is defined as the mean cosine of the scattering angle θ [22, 23], which describes the amount of directionality retained after a single scattering event such that *g* = 1 for entirely forward scattering, *g* = 0 for isotropic scattering, and *g* = -1 for entirely backward scattering. The diffusion approximation thus lumps together ***μ**_s_* and *g* [24].

More recently, optical coherence tomography (OCT) has been used to extract the optical properties of human tissues and optical phantoms [25, 26]. OCT is a non-invasive imaging technique that generates high resolution tomographic images using low coherent near infrared radiation (NIR). It measures characteristic patterns of directly elastically backscattered (low coherent ballistic and near-ballistic) photons from different reflective layers in a sample, e.g. at refractive index mismatches between tissue compartments with different microstructural properties [27]. If the OCT light source and imaging optics are calibrated with respect to absolute reflectivity, OCT can be used to obtain quantitative information on the scattering properties of the sample [25, 26, 28]. Levitz et al. [25] developed a theoretical model, based on inverse Monte Carlo modelling, to fit the depth dependent attenuation of the OCT signal and the local OCT signal intensity to extract ***μ**_s_* and *g*, respectively (see methods for details). If the investigated tissue/structure does not absorb strongly in the wavelength range of the light source, light attenuation in OCT is primarily a function of light scattering, where the local reflectivity quantifies the amount of directly backscattered photons, which can thus be used to quantify the *g* value [25] as a decrease in *g* causes an increase in the local reflectivity and *vice versa*. Another advantage of OCT is that it allows for very localised extraction of optical properties, while e.g. diffusion theory averages the optical properties over the measurement area (usually encompassing several cm^2^ of surface area) [21].

In the present study, we use OCT to characterise the optical scattering properties of intact coral tissues and skeletons *in vivo*. The specific aims are to study the variability in the scattering coefficient ***μ**_s_* and *g* value of 8 coral species with a specific focus on differences between tissue and skeletal scattering as well as differences among tissue types.

## Methods

### Coral specimens

The corals used in this experiment originated from the coral culture at the Centre Scientifique de Monaco. Corals were kept in aquaria supplied with Mediterranean seawater (exchange rate 70%/h) at a salinity of 38, temperature of 25°C, and photon irradiance (PAR, 400-700 nm) of 200 μmol photons m^−2^ s^−1^ on a 12-h/12-h photoperiod as provided by HQI-10000K metal halide lamps (BLV Nepturion). Corals were fed twice a week with *Artemia salina nauplii*. We selected several coral fragments from the following coral species: *Echinopora lamelosa, Platygyra pini, Hydnophora pilosa, Pavona cactus, Turbinaria reniformis, Acropora sp., Montipora capricornis* and *Stylophora pistillata*. The corals were chosen to represent a diversity of coral species with different tissue thicknesses, host pigmentation as well as skeletal optical properties [14]. At least three coral fragments of each species were used for optical extraction.

### Theory of optical parameter extraction based on OCT

The basic OCT operating principle is explained elsewhere [27]. OCT can be used to extract the scattering coefficient ***μ**_s_* and anisotropy of scattering, *g*, from biological tissues using a theoretical model of OCT light propagation derived from the inverse Monte Carlo method [25, 26, 28]. For a homogenous biological tissue, the depth dependent OCT reflectance signal, *R(z)*, is thereafter described as a simple exponential decay:

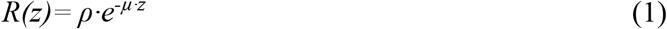

where *ρ* is the fraction of light sampled from the focal volume of tissue, and *μ* is the signal attenuation to and from the focal volume (see Levitz et al.[25] for details). Further,

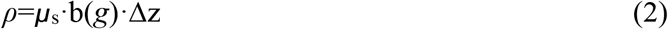

and

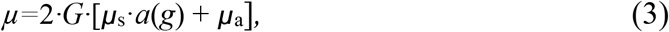

Here, Δ*z* is the axial resolution of the imaging system given by the coherence length *lc*:

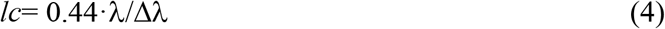

where *λ* is the center wavelength of the light source (930nm) and Δ*λ* is the spectral bandwidth (63.4 nm). The parameter b(*g*) is the scatter collection efficiency factor that describes the fraction of light scattered within the coherence gate (i.e. the spatial distance over which the reflected light and reference beam can cause interference signals) that is backscattered within the solid angle of collection by the objective lens. *G* is a geometry factor that accounts for the enhanced pathlength due to off-axis light propagation during delivery and collection by the objective lens. For an objective with low numerical aperture (NA), *G* is approximated by 1/cos[sin^−1^(NA)][25]. The theoretical model further assumes a Henyey-Greenstein scattering phase function *p*(θ) to describe the scattering of coral tissues, given that this phase function is most commonly used to describe the scattering of biological tissues [22, 29]. The phase function *p*(θ) affects the parameter b(*g*) by:

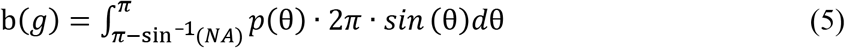

The factor *a* in Eq. 3 is the scattering efficiency factor that depends on *g*. It determines the ability of photons to reach the focus of the OCT system despite scattering; when *a* = 1 then *g* = 0 and *vice versa* [30]. The function *a*(*g*) can be approximated by Monte Carlo simulations [25] yielding:

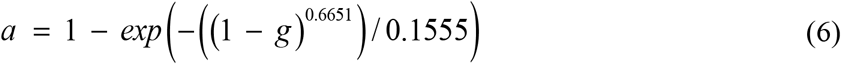

### OCT imaging and signal calibration

OCT imaging was performed on corals as described previously [31]. Briefly, imaging was done with a commercially available spectral-domain (SD) OCT system (Ganymede II, Thorlabs GmbH, Dachau, Germany) equipped with an objective lens with an effective focal length of 36 mm, and a working distance of 25.1 mm (LSM03; Thorlabs GmbH, Dachau, Germany; Fig. 1a). The system used a 930 nm light source, yielding a maximal axial and lateral resolution in water of 5.8 μm and 8 μm, respectively. OCT b-scans were acquired at a fixed pixel size of 581 × 1023, while the actual field of view was variable in y but fixed in z (z=2.2 mm). OCT imaging was performed for different coral species in a black acrylic flow chamber supplied with aerated seawater. For all measurements the OCT system was optimised to yield highest OCT signal measurements at a fixed distance, chosen to be in the upper 1/3 of the OCT image (at 0.4 mm from the top). To calibrate the OCT reflectivity, calibration measurements were performed prior to measurements for each coral species using a reflectance standard composing of (1) oil-glass interface, (2) water-glass interface, and (3) air-glass interface. The reflectivity from the reflectance standard was calibrated via Fresnel’s equations for the case of normal incidence of light using the refractive index of air (*n*=1), water (*n*=1.33), oil (*n*=1.46) and glass (*n*=1.52). The reflectance of the oil-glass interface was thus ((1.46-1.52)/(1.46+1.52))^2^ = 4.05×10^−4^, while the reflectance from a mirror = 1 (Fig. S1). The OCT signal from measurements on corals, OCT(dB), was converted to *R* via a linear fitting procedure, where the relation of OCT(dB) to log_10_(R) was described by linear function, as both parameters are logarithmic. To calibrate for the focus function of the objective lens, calibration measurements were performed for 8 different *z* positions (0.0 – 0.8 mm from the image top in steps of 0.1 mm). Such measurements showed that the OCT signal fall-off, from the z position of peak reflectivity (z= 0.4 mm) to z=0.8 mm, was exponential, facilitating a straight forward correction of acquired scans for the focus function of the OCT system.

**Fig. 1.**
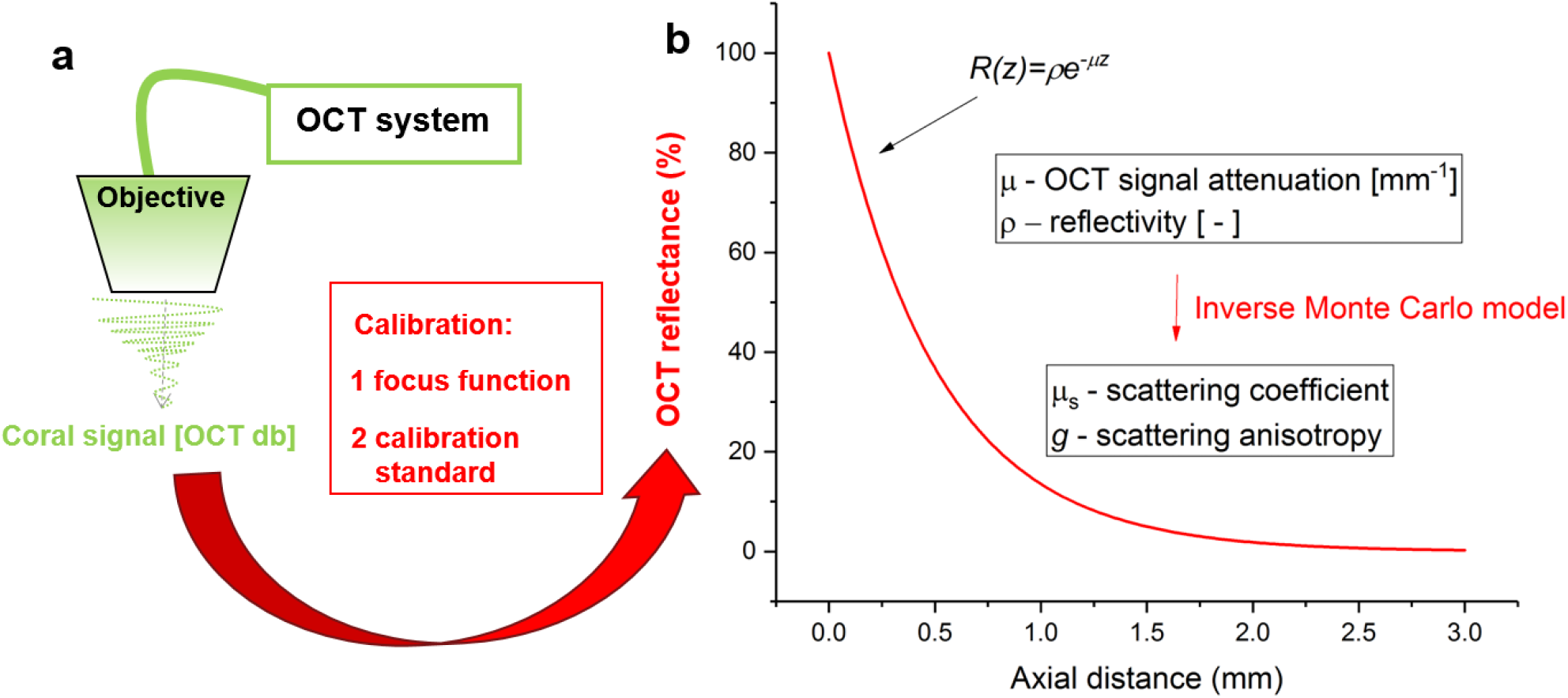
Experimental and theoretical approach to extract the scattering coefficient μs [mm-1] and anisotropy of scattering g [dimensionless] for intact living corals. a) A spectral domain OCT system provided non-invasive low coherent light (930 nm) that imaged corals in vivo without any actinic effect. For each coral, the raw OCT dB signal was calibrated via correcting for the depth-dependent signal attenuation (focus function calibration) and scanning against known reflectance standards to yield the absolute OCT reflectance. b) The vertical coral OCT reflectance attenuation at a given area of interest was then matched to a simple linear exponential function that yielded (1) the reflectivity and (2) the slope of the signal attenuation, which was then matched to gain μs and g via an inverse Monte Carlo approach (see methods for details).

### Optical extraction

After OCT scans were corrected for the focus function (OCT dB) and converted to absolute values of reflectivity (log_10_(R)) dedicated coral tissue areas were selected for optical extraction. Areas were selected to cover different tissue types, including polyp and coenosarc tissues as well as tissues covered with host pigment granules. Values of *ρ* (local reflectivity, dimensionless) and *μ* (linear signal attenuation [cm^−1^]) were matched to *g* and *μ_s_* using the theory described above [25, 30]. The absorption coefficient, ***μ**_a_*, by coral and algal pigments at 930 nm is negligible [14, 16] and the fitting assumed that absorption was dominated by water, where *μ_a_*= 0.43 cm^−1^ for water at 930 nm [22]. The water content of the tissue was assumed to be 50%, based on the hydration of human tissues [22]. The effective numerical aperture was 0.15 and the OCT coherence length was 6 μm. Based on these settings a grid method was applied [25] to extract the optical parameters *g* and *μ_s_*. Briefly, the grid method is a lookup table of experimental values of *μ* versus *ρ* generated by choosing values of *μ_s_* and *g* for use in Monte Carlo simulations to create *R(z)* curves that are fit using Eq. 1 to yield *μ*(*μ_s_,g*) and *ρ*(*μ_s_,g*) (Fig. 1). When the *μ*(*μ_s_,g*) and *ρ*(*μ_s_,g*) values are plotted, and the iso-*μ_s_* and iso-*g* lines are drawn, a grid is formed. A pair of observed *μ,ρ* values now uniquely maps to a pair of *μ_s_,g* values [25]. A 2D interpolation algorithm implements the mapping, specifying *μ_s_*(*μ,ρ*) and *g*(*μ,ρ*). More details on the grid can be found in [25, 30].

Optical extraction involved curve fitting for randomly selected tissue and skeletal spots. Curve fitting was considered satisfactory if the model matched the data with a R^2^ value >0.5. The first few pixels at the tissue surface were excluded from the curve fitting procedure [32], given the strong reflectivity spike due to the refractive index mismatch between tissue and water. Preliminary analyses showed that the OCT signal often did not attenuate within the first 20-50 μm of the coral tissue (Fig. 2). Maximal OCT signal was found at about 50 μm depth and curve fitting used maximal reflectivity values that showed a smooth signal fall off. We found that optical extraction for regions of interest positioned deeper within the tissue, e.g. underlying coral skeletons, would not yield reasonable results due to OCT signal attenuation by upper tissue layers. More robust estimates of coral skeleton optical properties were obtained from OCT scans of bare skeletons, where the tissue was removed with an air gun [15].

**Fig. 2.**
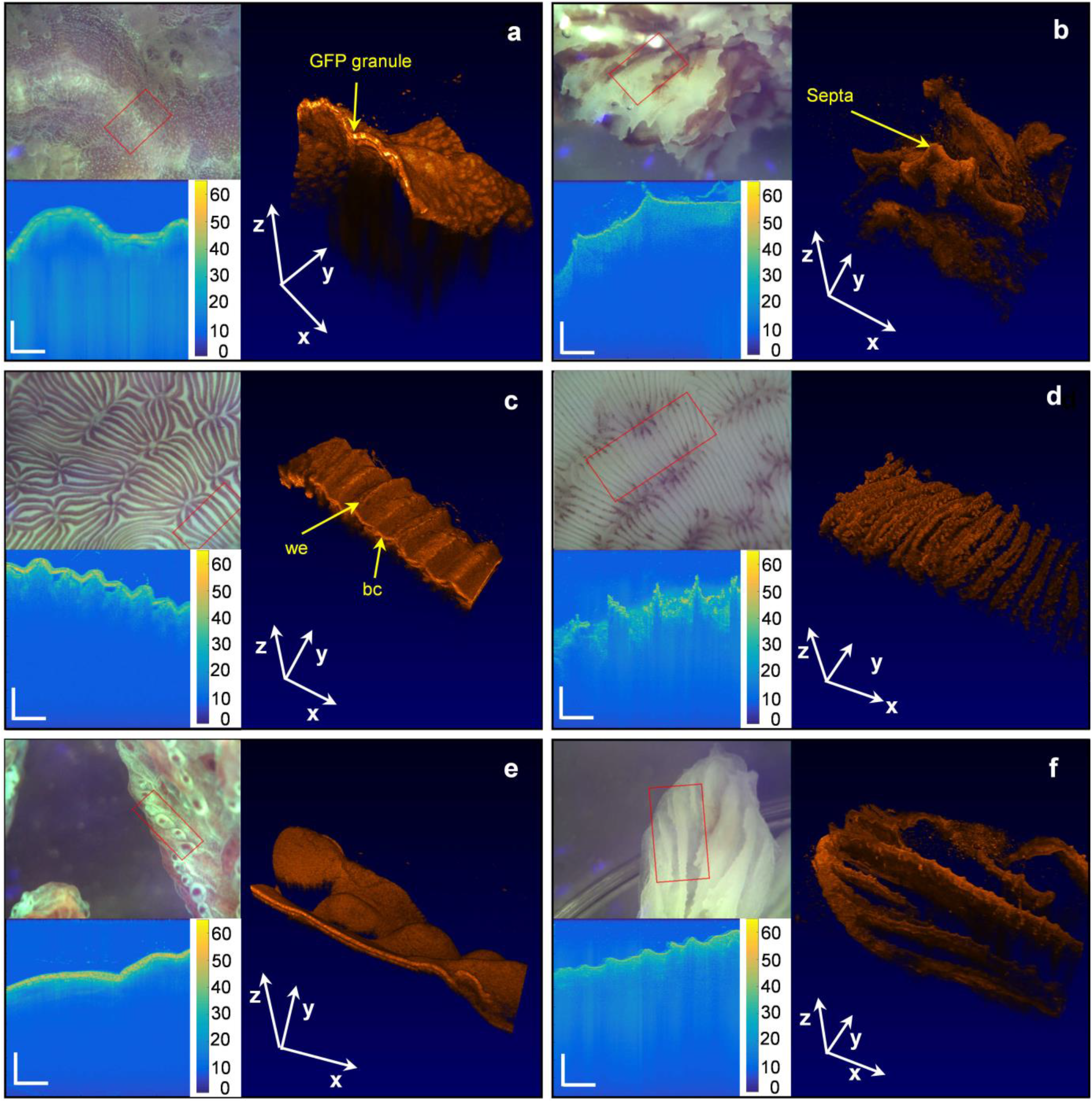
Overview of OCT scanning on intact corals (left panel) and bare coral skeletons (right panel). OCT scans are shown for *Platygyra pini* (**a, b**), *Pavona cactus* (**c, d**) and *Hydnophora pilosa* (**e, f**). Close-up photographs were taken with the USB camera of the OCT system. The area within the red square corresponds to the three-dimensional OCT scans (displayed in black-orange false-color coding, which was adjusted for optimised visualisation). The *x-y-z* scale bars represent a distance of 2 mm in each dimension. Exemplary cross-sectional OCT B-scans are shown before calibration with the false-color legend representing the OCT signal from 0-60 OCT dB. Scale bars in z and x dimensions represent a distance of 400 μm. we= white elevation, bc= brown crevices.

## Results and discussion

### Light scattering in tissue vs skeleton

The inherent optical properties of coral tissues are largely unknown, although they are fundamental for a better mechanistic understanding of coral light absorption and photosynthesis, and thus coral ecophysiology [16, 33, 34]. Previous studies have primarily characterised the apparent optical properties of coral tissues, i.e., light field parameters such as the scalar irradiance and reflectance, suggesting that coral tissues are highly scattering [15, 35]. However, only one study has presented data on the scattering coefficient of intact coral tissues, providing evidence for a high reduced scattering coefficient, *μ_s_*’, of a massive faviid coral tissue [16]. In the present study, determination of *μ_s_* in intact coral tissue for 8 corals showed high variability of *μ_s_* from 4 to 25 mm^−1^ at 930 nm (Table 1). In comparison, *μ_s_* of plant tissues ranges from about 5 to 10 mm^−1^ [36, 37], while human skin can reach values of about 20 mm^−1^ [22]. Thus for the case of a highly scattering coral tissue, such as in *Platygyra pini* (Fig. 2), the average distance between scattering events, i.e, the scattering mean free path, MFP=1/μ_s_, is 46 μm (Table 1), which is about 2-fold lower than for most biological tissues (MFP~100 μm) [38]. In contrast, the scattering strength of the coral *Stylophora pistillata* would rank among the lower end of biological tissues, with a MFP of about 250 μm (Table 1) [22].

**Table 1.** Optical properties of 8 coral species extracted from calibrated OCT scans at 930 nm. The extracted *ρ* (local reflectivity) and μ (signal attenuation) were used to fit g (anisotropy of scattering) and *μ_s_* (scattering coefficient). The mean free path (MFP=1/ *μ_s_*) and the transport mean free path (TMFP=1/ *μ_s_*’) were calculated. Data are means (±SE).

| Species | Type | $\rho$<br>[] | $\mu$<br>(mm<sup>-1</sup>) | $g$<br>[] | $\mu_s$<br>(mm<sup>-1</sup>) | $MFP$<br>(mm) | $TMFP$<br>(mm) | $R^2$ | $n$ |
| --- | --- | --- | --- | --- | --- | --- | --- | --- | --- |
| *Acropora sp.* | tissue | 2.10E-06<br>(±4.04E-07) | 10.5<br>(±1.1) | 0.98<br>(±0.003) | 14.5<br>(±1.44) | 0.08<br>(±0.01) | 5.34<br>(±0.82) | 0.56<br>(±0.03) | 10 |
| *Echinopora lamelosa* | tissue | 5.45E-06<br>(±7.85E-07) | 20.9<br>(±1.5) | 0.97<br>(±0.005) | 25.2<br>(±1.8) | 0.04<br>(±0.001) | 2.1<br>(±0.29) | 0.79<br>(±0.01) | 22 |
| *Hydnophora pilosa* | tissue | 4.64E-06<br>(±7.93E-07) | 13<br>(±1.2) | 0.96<br>(±0.007) | 12.2<br>(±1.09) | 0.09<br>(±0.01) | 2.7<br>(±0.44) | 0.71<br>(±0.04) | 12 |
| | skeleton | 8.19E-07<br>(±1.60E-07) | 7.7<br>(±0.85) | 0.99<br>(±0.003) | 12.1<br>(±1.38) | 0.09<br>(±0.01) | 7.34<br>(±0.76) | 0.55<br>(±0.03) | 6 |
| *Montipora capricornis* | tissue | 6.72E-05<br>(±2.04E-05) | 16.2<br>(±1.6) | 0.59<br>(±0.09) | 8.63<br>(±0.83) | 0.12<br>(±0.01) | 0.62<br>(±0.31) | 0.75<br>(±0.03) | 8 |
| *Pavona cactus* | tissue | 1.75E-05<br>(±9.13E-06) | 20.5<br>(±2.3) | 0.91<br>(±0.03) | 17.4<br>(±1.65) | 0.06<br>(±0.01) | 1.53<br>(±0.31) | 0.78<br>(±0.02) | 14 |
| | skeleton | 2.12E-06<br>(±4.96E-07) | 10.8<br>(±1.2) | 0.98<br>(±0.004) | 15.6<br>(±1.8) | 0.07<br>(±0.009) | 0.52<br>(±0.13) | 0.56<br>(±0.03) | 6 |
| *Platygyra pini* | tissue | 1.42E-4<br>(±7.51E-05) | 26<br>(±3.6) | 0.69<br>(±0.11) | 21.3<br>(±2.02) | 0.05<br>(±0.01) | 1.37<br>(±0.45) | 0.79<br>(±0.03) | 11 |
| | skeleton | 1.04E-04<br>(±1.96E-05) | 6.0<br>(±0.74) | 0.15<br>(±0.05) | 2.7<br>(±0.35) | 0.44<br>(±0.05) | 0.55<br>(±0.08) | 0.54<br>(±0.01) | 12 |
| *Stylophora pistillata* | tissue | 3.55E-05<br>(±1.74E-05) | 6.3<br>(±0.5) | 0.51<br>(±0.16) | 4<br>(±1) | 0.33<br>(±0.06) | 2.13<br>(±1.01) | 0.81<br>(±0.03) | 6 |
| | skeleton | 7.59E-05<br>(±2.59E-05) | 8.4<br>(±1.4) | 0.28<br>(±0.05) | 4.03<br>(±0.74) | 0.36<br>(±0.09) | 0.52<br>(±0.13) | 0.53<br>(±0.04) | 9 |
| *Turbinaria reniformis* | tissue | 4.69E-06<br>(±1.01E-06) | 15<br>(±2) | 0.96<br>(±0.02) | 17.3<br>(±3.26) | 0.08<br>(±0.01) | 2.32<br>(±0.29) | 0.77<br>(±0.03) | 9 |

The relative role of tissue vs skeleton light scattering in modulating light propagation for coral photosynthesis has been debated [16, 18, 19]. We thus compared tissue and skeletal scattering on the same coral colonies using OCT *in vivo* (Fig. 2, Table 1). A comparison of tissue and skeletal scattering showed that tissue scattering was 8-fold higher compared to skeleton scattering for *P. pini* (Table 1), supporting earlier findings of high tissue vs low skeleton scattering in a comparable faviid coral [16]. In contrast, for the corals *S. pistillata, P. cactus* and *H. pilosa, μ_s_* was similar between tissue and skeletons (Table 1). These results highlight species-specific variations in the relative role of tissue and skeletal light scattering for coral light management.

### Light scattering characteristics of different tissue types

The structural organisation of coral tissues varies within a coral species [31], and given that light scattering in biological tissues is a function of refractive index fluctuations due to different cell types and constituents [22], it is thus to be expected that scattering variability exits within different spatial compartments of the coral tissue. We found that the scattering coefficient of coenosarc tissues (*μ_s_*=24.6 ± 2.8 SE) was about 3-fold higher compared to polyp tentacle tissue (*μ_s_*=8.3 ± 0.4 SE) in the coral *Turbinaria reniformis* (one-way ANOVA F_1,7_= 26.3, p=<0.01, Fig. 3a-e). The lower scattering for the polyp tentacles could be related to a simpler tissue structure as the tentacles do not have a mesoglea and an aboral gastrodermal layer as compared to the coenosarc tissue [39]. The mesoglea is collagen-rich and has been suggested to have an important role in tissue light scattering [15, 16, 40].

Close up images of the tissue surface of the coral *Pavona cactus* revealed an interesting tissue surface pattern, alternating between ‘brown crevices’ and ‘white elevations’ (Fig, 2c, 3f). Although we did not characterise *Symbiodinium* distribution via spectral measurements, the brown colour is clearly indicative of a dense aggregation of *Symbiodinium* cells [41]. Cross-sectional OCT scanning showed that ‘white elevations’ are skeletal extrusions, fully covered by living tissue but apparently lacking *Symbiodinium* (Fig. 3g). These extrusions are likely coenosteal spines, which are found in branching corals, including e.g. *Stylophora pistillata* and *Pocillopora damicornis* [42, 43]. Optical analyses showed that the scattering coefficient was about 1.8-fold higher for the brown crevices (*μ_s_*= 22.42 ± 1.67 SE) compared to the white elevations (*μ_s_*= 12.35 ± 0.76 SE) (one-way ANOVA F_1,12_= 29.9, p=<0.01, Figure 3). The enhanced scattering found in the brown crevices compared to the skeletal elevations of the coenosarc tissue of *Pavona cactu*s could indicate a link to the aggregation of *Symbiodinium* cells. The scattering coefficient of *Symbiodinium* cells was estimated to *μ_s_*=16 mm^−1^ [20], which is 1.3 fold higher than the average scattering over the white areas (Fig. 3h). The scattering of *Symbiodinium* likely depends on the clade specific cell ultrastructure [44], including e.g. the density of membrane lipids, which are important light scatterers [45, 46]. Although a detailed study on the scattering coefficient of *Symbidinium* was beyond the scope of the present study, these results hint towards a potential role of light scattering by *Symbiodinium* for coral light management, which warrants future investigations.

**Fig. 3.**
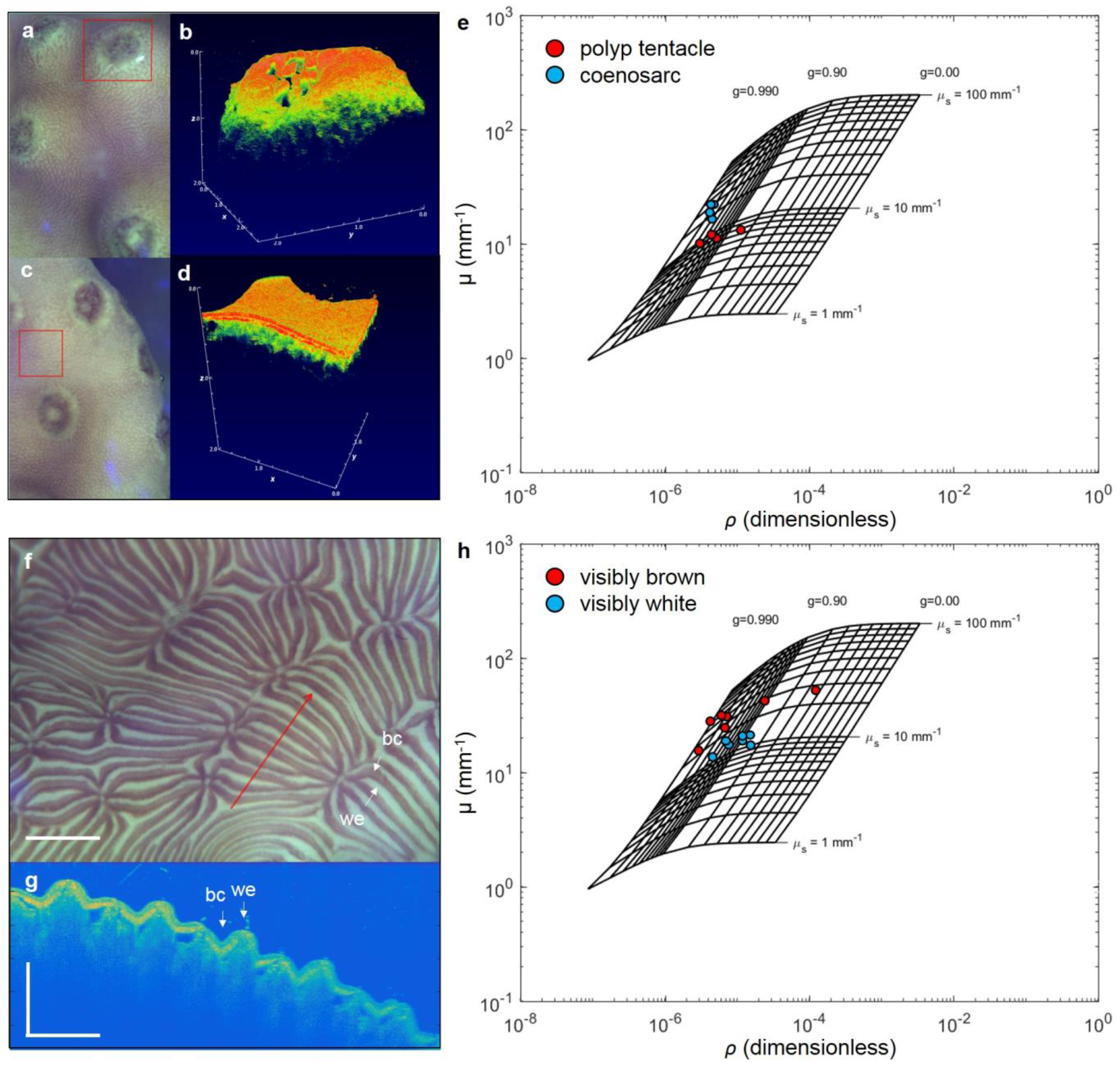
Characterisation of scattering properties of different coral tissue types. Close up images (**a,c**) and cross sectional OCT B-scans (**b,d**) of *Turbinaria reniformis* coenosarc and polyp tissues, respectively (as shown in red square). The optical grid (**e**) matches the local reflectivity (*ρ*) to the anisotropy of scattering, *g*, while the vertical OCT signal attenuation,*μ*, yields the optical scattering coefficient, *μ_s_*. (see methods for details). Close-up image (**f**) of the tissue surface of *Pavona cactus* covering brown cervices (bc) and white elevations (we). Scale bar = 2 mm. The red arrow shows the area covered by OCT imaging in panel b. Exemplary OCT cross-sectional B-scan (**g**). Scale bar in *x* and *z* = 500 μm. Optical grid comparing white elevations and brown crevices (**h**).

We also studied the light scattering properties of fluorescent host pigments (Fig. 4a-g). The role of fluorescent host pigments (FP) for coral photosynthesis and ecophysiology is disputed, as studies have shown that FP can be either photoprotective (e.g. by absorbing damaging UV radiation and backscattering of incident radiation [47]) or stimulate photosynthesis [48–50] depending on their spectral properties and distribution in the coral tissue. It has also been suggested that such photoprotective and/or photosynthesis stimulating functions depend on the type of FP and its structural aggregation within the tissue [31, 51]. In our study, we found FP aggregations in *Platygyra pini*, where they formed cluster-like ~50-100 μm wide granules also known as chromatophores [31, 49] (Fig. 2a, Fig. 4e,f). These chromatophores are composed of smaller scattering granules, typically about 1 μm in diameter [49]. Our optical analyses showed that tissue areas with such FP aggregates have a low *g* value of about 0.34 (± 0.09 SE) indicating a nearly isotropic scattering behaviour of FP granules (Fig. 4g). In contrast, coral tissue with a more diffuse distribution of FP was strongly forward scattering with a g value of about 0.96 (± 0.008 SE, pooled for *Hydnopora pilosa* and *Echnipora lamelosa*) (Fig. 4a-d,g). Likewise, brown-pigmented tissue in *P. pini* was forward scattering (g=0.97 ± 0.004 SE (Fig. 4g).

**Fig. 4.**
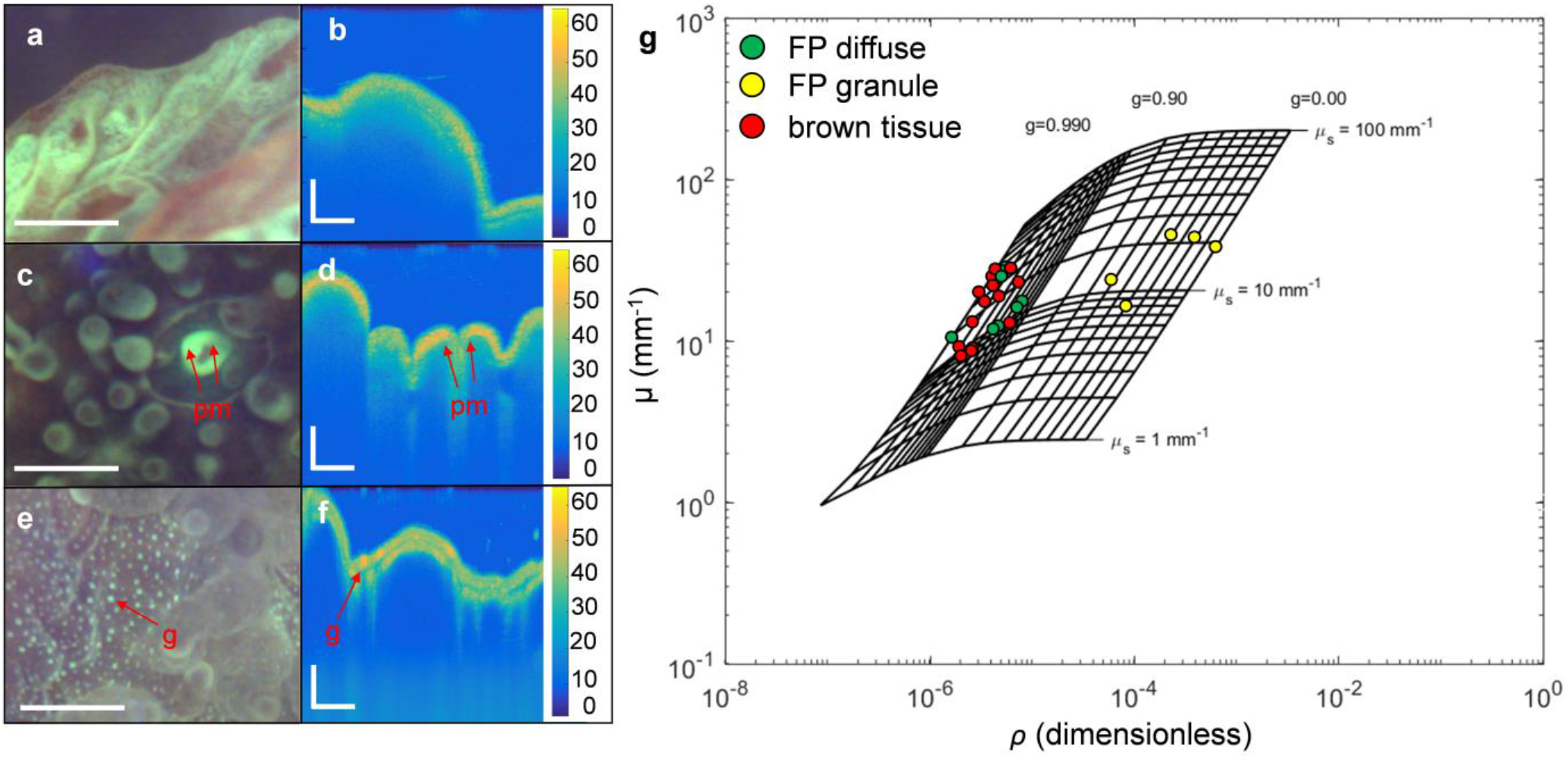
Light scattering by fluorescent host pigments (FP) in *Hydnophora pilosa* (a,b), *Echinopora lamelosa* (c,d) and *Platygyra pini* (e,f). pm= polyp mouth, g= granule. The optical analyses **(g)** included all three species and show extracted *g* values for brown tissue areas without visible FP content, tissue areas with a diffuse FP distribution, and tissue areas with a clear aggregation of FP into granules. Scale bars are 1mm (a, c, e) and 200 μm (b, d, f).

Previous studies showed that high densities of light scattering FP granules lead to enhanced tissue reflectivity and surface scalar irradiance [11, 51]. Many faviid corals have a dense network of GFP granules, covering >30% of the polyp surface area [31]. Our results suggest that the almost isotropic scattering behaviour of FP granules leads to an effective lateral homogenization of the vertically incident irradiance, which is characterised by enhanced tissue surface scalar irradiance and a more rapid vertical attenuation of irradiance compared to tissue areas with diffuse FP [11]. The *g* value is diagnostic for the size of the scattering elements and assuming that the light scattering particles are spherical, a lower *g* value hints towards a smaller size of the scattering elements [52]. Certainly, links between ultrastructure, scattering properties of FP granules, and the photobiology of corals warrant further investigation.

### Implications of coral scattering properties for coral light management

The present study provides detailed *in vivo* estimates of the inherent optical properties of live corals which is a key requirement for a better mechanistic understanding of coral optics and thus coral photosynthesis [13]. The quantification of light scattering (***μ**s* and *g*) allows for improved light propagation models [16, 20]. Although a detailed simulation was beyond the scope of the present study, one can exemplify basic principles of coral light harvesting in a simplified simulation (see Supplementary Information). We developed a Monte Carlo simulation for two corals (*Stylophora pistillata* and *Hydnophora pilosa*) with different light scattering properties (Table, Fig. S2). Previous studies suggested that the optical properties of the skeleton of *S. pistillata* lead to a strong lateral redistribution of incident irradiance which enhances photosynthetic efficiency but also makes this species highly susceptible to coral bleaching [14, 18].

Using the *in vivo* optical properties of coral tissue and skeleton (Table 1) we show that incident light is indeed strongly redistributed by light scattering in *Stylophora pistillata*. The high lateral spread of light leads to an 84% absorption of light by the tissue layer (assuming ***μ**a*= 3 mm^−1^, Fig. S2a). In contrast, the light scattering properties of *H. pilosa* lead to a much lower lateral spread of light and an approximate 1.3-fold reduced light absorption by the tissue layer (Fig. S2b). Although the scattering coefficient in *H. pilosa* is higher than in *S. pistillata* (Table 1), the *g* value in *S. pistilita* is very low (tissue, g=0.51 skeleton, g=0.28) leading to a more isotropic scattering behaviour and enhanced lateral spread of light (Fig. S2a). Most biological tissues are strongly forward scattering [22] and *g* values of 0.9 have been assumed for corals [16]. However, our study shows that the assumption of strongly forward scattering tissues is not always correct for coral tissues and skeletons (Table1).

It is important to point out, that coral morphology (mm to cm scale) also affects light distribution [18, 53] and first attempts have been made to model light propagation with a 3D architecture [16]. Mesh-based 3D monte Carlo simulations have been developed for brain tissues in optogenetics [54] and can now in theory also be developed using 3D segmented tissue and skeletal architectures from OCT scans (Fig. 1) [31] combined with data on scattering properties of intact corals (this study) and skeletons [14, 20].

In conclusion, the current OCT-based approach allows for a rapid determination of *in vivo* optical properties of corals *in vivo* under defined laboratory conditions. The experimental measurements are fast and application of the theoretical model is straight forward, showcasing the suitability of OCT as a rapid method for optical characterisation. Our approach is currently limited to the extraction of optical properties in the NIR range (930 nm) and ideally this should be improved for characterising light scattering in the visible range (400-700nm) via e.g. visible light OCT [55] or diffuse optical spectroscopy [22].

## Author Contributions

DW, CFP and MK designed study. DW, JBH and NL performed measurements. SLJ provided analytical tools. CFP provided coral holding and lab facilities, maintained and prepared coral specimens. DW analyzed data with input from SLJ and MK. DW wrote the article with editorial input from all co-authors.

## Acknowledgments

We acknowledge Cecile Rottier and other staff at the Centre Scientifique de Monaco for help and excellent technical assistance.

